# A triple-hybrid cross reveals a new hybrid incompatibility locus between *D. melanogaster and D. sechellia*

**DOI:** 10.1101/590588

**Authors:** Jacob C. Cooper, Ping Guo, Jackson Bladen, Nitin Phadnis

**Affiliations:** School of Biological Sciences, University of Utah, Salt Lake City, UT 84112, USA

## Abstract

Hybrid incompatibilities are the result of deleterious interactions between diverged genes in the progeny of two species. In *Drosophila*, crosses between female *D. melanogaster* and males from the *D. simulans* clade (*D. simulans*, *D. mauritiana*, *D. sechellia*) fail to produce hybrid F1 males. When attempting to rescue hybrid F1 males by depleting the incompatible allele of a previously identified hybrid incompatibility gene, we observed robust rescue in crosses of *D. melanogaster* to *D. simulans* or *D. mauritiana*, but no rescue in crosses to *D. sechellia*. To investigate the genetic basis of *D. sechellia* resistance to hybrid rescue, we designed a triple-hybrid cross to generate recombinant *D. sechellia* / *D. simulans* genotypes. We tested the ability of those genotypes to rescue hybrid males with *D. melanogaster*, and used whole genome sequencing to measure the *D. sechellia* / *D. simulans* allele frequency of viable F1 males. We found that recombinant genotypes were rescued when they contained two specific loci from *D. simulans* – a region containing previously identified *Lethal hybrid rescue (Lhr)*, and an unknown region of chromosome 3L which we name *Sechellia aversion to hybrid rescue (Satyr)*. Our results show that the genetic basis for the recent evolution of this hybrid incompatibility is simple rather than a highly dispersed effect. Further, these data suggest that fixation of differences at *Lhr* after the split of the *D. simulans* clade strengthened the hybrid incompatibility between *D. sechellia* and *D. melanogaster*.

## Introduction

*Drosophila melanogaster* and its sister species in the *Drosophila simulans* clade are a classic model for studying hybrid incompatibilities. With a rich history spanning more than 100 years, the problem of understanding the genetic basis of hybrid lethality in this system has been under consistent, creative, and often surprising lines of attack (Barbash 2010). Through a combination of X-ray and chemical mutagenesis and the isolation of natural rescue alleles, three hybrid incompatibility genes required for F1 male lethality between *D. melanogaster* and *D. simulans* have been identified so far– *Hybrid male rescue* (*Hmr*), *Lethal hybrid rescue* (*Lhr*), and *GST-containing FLYWCH Zinc-Finger protein* (*gfzf*) (Pontecorvo 1943; Watanabe 1979; Hutter and Ashburner 1987; Barbash *et al.* 2003; Brideau *et al.* 2006; Phadnis *et al.* 2015). In hybrids, only one allele of each gene is incompatible (*Hmr*^*mel*^, *Lhr*^*sim*^, and *gfzf*^*sim*^) (Figure 1B). Loss of any of the incompatible alleles rescues the viability of hybrid F1 males. *Hmr* and *Lhr* are heterochromatin associated transcriptional repressors that physically bind each other and suppress the expression of transposable elements and repetitive DNA sequences (Thomae *et al.* 2013; Satyaki *et al.* 2014). *gfzf* is a general transcriptional co-activator for TATA-less genes along with M1BP (Baumann *et al.* 2018). Recent work also demonstrates that *Hmr* mislocalizes to *gfzf* bound chromatin in F1 hybrids, indicating that they have the capability to interact with many other loci (Cooper *et al.* 2018). Though the identity of these three genes has been established, there is still no comprehensive explanation of their role in hybrid incompatibility, and an incomplete picture of other genes they interact with to cause hybrid incompatibility.

**Figure 1.**
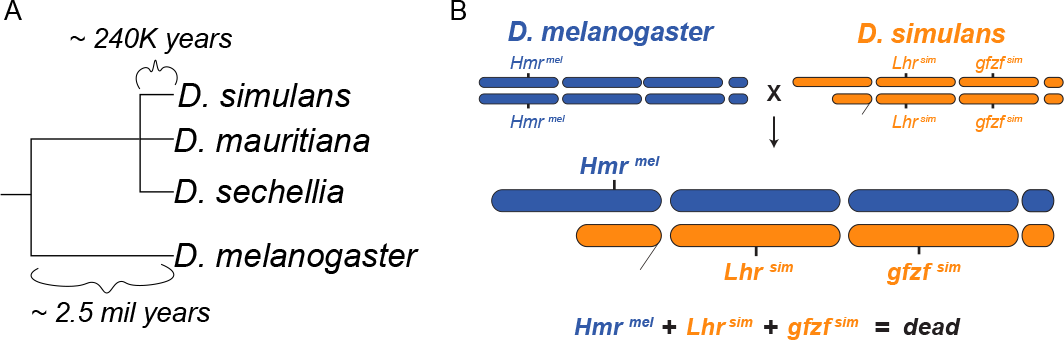
Hybrid incompatibilities between *D. melanogaster* and the *D. simulans* clade. (A) Cladogram for the *D. melanogaster* – *D. simulans* clade relationship. (B) Schematic of the hybrid incompatibility genes between *D. melanogaster* and *D. simulans*

The *Drosophila simulans* clade contains three species (*D. simulans, D. mauritiana*, and *D.sechellia*) which diverged from their last common ancestor around 240 thousand years ago (Garrigan *et al.* 2012) (Figure 1A). *D. melanogaster* has strong F1 hybrid incompatibilities with all members of the *D. simulans* clade, which all follow a common pattern – in particular, *D. melanogaster* females crossed with *D. simulans* clade males do not produce F1 hybrid males and produce sterile F1 hybrid females (Sturtevant 1919). The genetic architecture of the hybrid incompatibility between *D. melanogaster* and the other species of the *D. simulans* clade have shared elements, and the genetic architecture of hybrid incompatibilities have shared features. Alleles of *Hmr* that rescue hybrid F1 males between *D. melanogaster - D. simulans* also rescue hybrid F1 males between *D. melanogaster - D. mauritiana* and *D. melanogaster* – *D. sechellia* (Hutter and Ashburner 1987). However, hybrid male rescue between *D. melanogaster* – *D. sechellia* occurs at lower frequency than the other hybridizations. *D. sechellia* is an island species that is specialized on the toxic Morinda fruit (Tsacas 1981). Though they have some continued introgression with *D. simulans* (Matute and Ayroles 2014), they are largely isolated from *D. simulans* by extensive ecological diversification (Huang and Erezyilmaz 2015) and F1 male sterility (Lachaise *et al.* 1986). However, there has been no attempt to account for the reduction in hybrid male rescue with *D. melanogaster*.

As such, the genetic basis for the lower rate of hybrid male rescue between *D. melanogaster* and *D. sechellia* remains unknown. Any resistance to hybrid male rescue alleles must act dominantly, since the equivalent *D. melanogaster* alleles are present in all F1 hybrids. It is clear that hybrid incompatibilities accumulate within the *D. simulans* clade (Cattani and Presgraves 2009; Garrigan *et al.* 2014) and that many recessive incompatibilities have accumulated between *D. melanogaster* and *D. simulans* (Presgraves 2003). Additionally, hybrid incompatibilities might evolve by many small gradual steps, distributing quantitative changes in hybrid rescue over many loci in the genome. This has proven to be the case with *Maternal hybrid recue*, a component of the female embryonic lethality observed in crosses between *D. melanogaster* males and female *D. simulans* females (Gérard and Presgraves 2012). The molecular properties of *Hmr*, *Lhr*, and *gfzf* indicate that they all have the capacity to interact with large fractions of the genome. Several proposals for their role in hybrid incompatibilities include: acting to buffer genome wide properties of chromatin or transposable elements (Brideau *et al.* 2006), general buffers against lethality (Castillo and Barbash 2017), or many moderate effects on different phases of the cell cycle (Cooper and Phadnis 2016).

Here, we use a triple-hybrid cross to dissect the interspecies variation in hybrid rescue between *D. melanogaster* and the *D. simulans* clade. We use knockdown of the *D. simulans* sibling species allele of *gfzf* (*gfzf*^*sib*^) to rescue hybrid males between *D. melanogaster* and the three species of the *D. simulans* clade. Surprisingly, we find that there is no rescue of hybrid males between *D. melanogaster* and *D. sechellia.* We design a cross to leverage the variation in rescue between *D. simulans* and *D. sechellia*, and generate a one generation recombinant QTL map of hybrid male rescue with pooled whole genome sequencing. We find that the lack of male rescue in *D. sechellia* is likely due to just two dominant major effect loci. One of these loci maps to chromosome 2R at the same genomic coordinates of *Lhr*. The other maps to a previously unidentified hybrid rescue locus on chromosome 3L, which we name *Sechellia aversion to hybrid rescue (Satyr)*. Our data suggests that the variation in hybrid rescue with respect to *gfzf* between *D. simulans* and *D. sechellia* is due to few changes of large effect size, and indicates that major components of *D. melanogaster* – *D. simulans* clade hybrid incompatibly remain unidentified.

## Results

### Knockdown of gfzf rescues hybrid males from crosses with D. simulans and D mauritiana, but not with D. sechellia

The closest sister species of *D. melanogaster* are the simulans clade, which includes *D. simulans*, *D. mauritiana* and *D. sechellia*. These three species are estimated to have diverged from their last common ancestor approximately 240,000 years ago, meaning they are relatively young species for *Drosophila* (Garrigan *et al.* 2012). The pattern of divergence of these three species is complex, and the phylogenetic relationship between these three species remains an unresolved trichotomy (Garrigan *et al.* 2012). *D. melanogaster* females carrying null mutations at *Hmr* produce viable males in crosses with males from any of the three sister species, indicating that the genetic basis of hybrid F1 male lethality between *D. melanogaster* and sister species is shared. Similar hybrid rescue crosses using null mutants of *Lhr* in *D. mauritiana* and *D. sechellia* are not yet possible, because hybrid rescue mutations in *Lhr* have only been isolated in *D. simulans*. As *gfzf*^*sim*^ is necessary for the lethality of hybrid F1 males in crosses between *D. melanogaster* females and *D. simulans* males, RNAi induced knockdown of *gfzf*^*sim*^ is sufficient to rescue the viability of hybrid F1 males (Phadnis *et al.* 2015). In these crosses, all necessary transgenes for produce hybrid rescue come from *D. melanogaster*, and the RNAi target sequences for *gfzf* are shared across all sibling species of the simulans clade (*gfzf*^*sib*^). This tool opens the door to test whether knockdowns of the *gfzf* allele from *D. mauritana* and *D. sechellia* are also sufficient to rescue hybrid male viability.

To assay for variation in hybrid male rescue by *gfzf*^*sib*^ knockdown across the *D. simulans* clade, we crossed *D. melanogaster* females carrying the *gfzf* knockdown constructs to males from several lines of *D. simulans*, *D. mauritiana*, and *D. sechellia*. In particular, we used two RNAi constructs that specifically target *gfzf* from the sister species at different regions of the gene. We first sequenced the RNAi target site from all our lines of *D. simulans*, and found perfect match for the target sequence in all of these strains. Similarly, the *D. mauritiana* strains are perfectly matched for the knockdown constructs with the exception of one line *w140*, which carries a single nucleotide mismatch for both knockdown constructs. *D. sechellia* is fixed for this single nucleotide mismatch for the first knockdown construct, but is perfectly matched for the second construct (Figure 2A).

**Figure 2.**
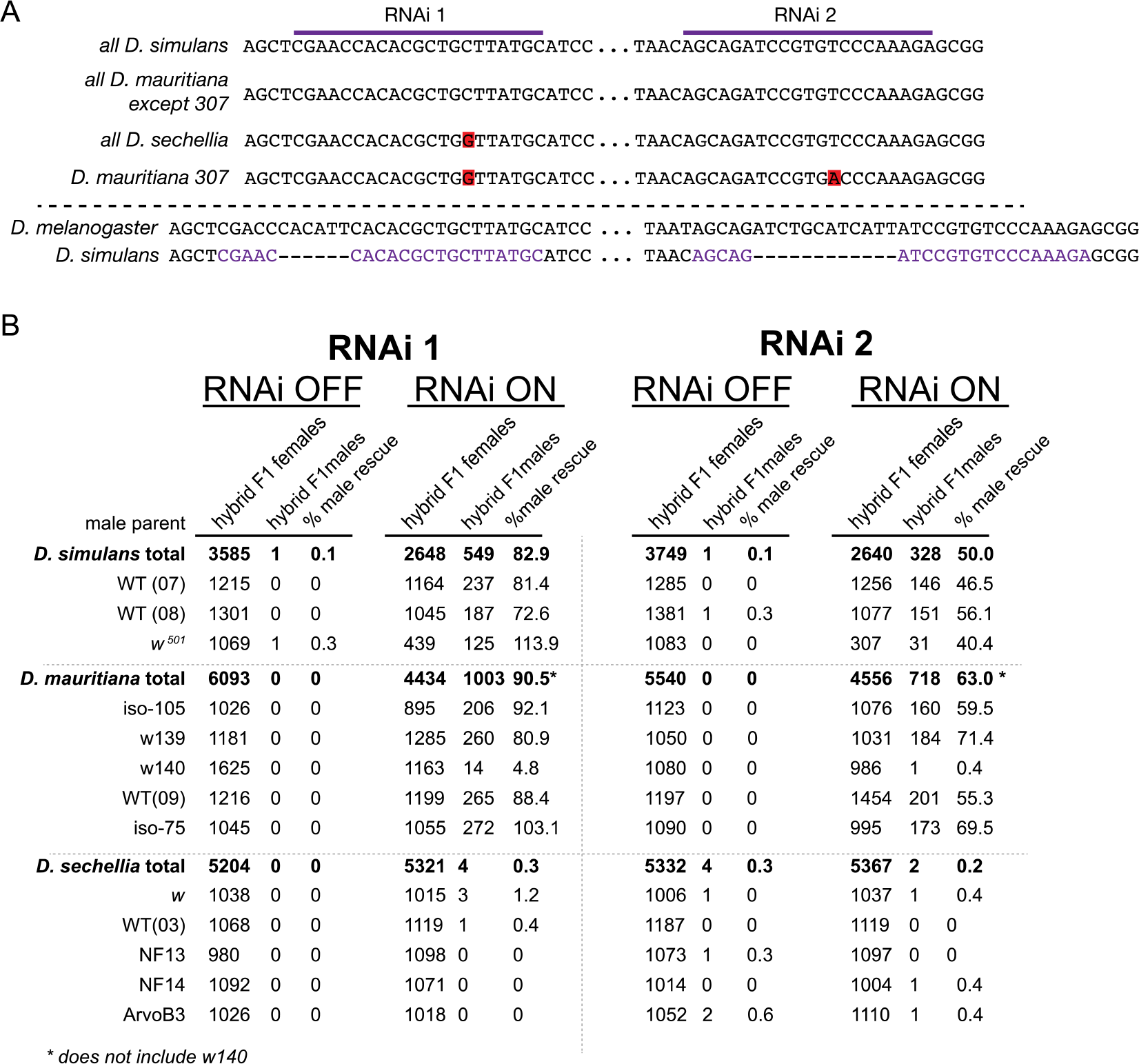
*gfzf* knockdown rescues hybrids with *D. simulans* and *D. mauritiana* but not *D. sechellia*. (A) Rescue crosses for hybrids with both *gfzf*^*sib*^ RNAi constructs. Summaries are presented for each species. (B) Alignment of RNAi targeting sites in all three *D. simulans* clade species. Below is an alignment with *D. melanogaster*, demonstrating the deletion that is fixed in the entire *D. simulans* clade.

In crosses with *D. melanogaster* females carrying *gfzf*^*sib*^ knockdown constructs, hybrid males from all lines of *D. simulans* showed robust rescue of hybrid male viability (Figure 2B). In similar *gfzf*^*sib*^ knockdown crosses with males from *D. mauritiana* strains, we observed robust rescue of hybrid F1 male viability at rates comparable or slightly better than those observed with *D. simulans*. Only one *D. mauritiana* line (*w140*) recorded no rescue, which may be explained by the mismatches for both knockdown constructs seen in this strain. Our results from crosses with *D. sechellia*, however, were dramatically different. In contrast to our observations of robust hybrid male rescue with *D. simulans* and *D. mauritiana*, we observed no rescue with *D. sechellia* with either RNAi construct. Although we did not sequence rare survivor males from these crosses, these are known to be the result of fertilization between nullo-X eggs from non-disjunction events in *D. melanogaster* and sperm carrying an X chromosome. Together, our results show that, unlike *Hmr*-based hybrid males rescue, RNAi targeting of the incompatible allele of *gfzf* is sufficient to rescue hybrid F1 male viability in crosses with *D. simulans* and *D. mauritiana*, but not with *D. sechellia*.

*D. sechellia* resistance to hybrid male rescue by *gfzf* knockdown may be explained by a failure to knockdown of *gfzf*^*sec*^ in *D. melanogaster*-*D. sechellia* hybrids. Because the short interfering RNA (siRNA) pathway genes such as Dicer-2, Ago-2, and R2D2, are rapidly evolving between *D. melanogaster* and the *D. simulans* clade, the siRNA pathway itself may not be as effective in these in *D. melanogaster*-*D. sechellia* hybrids (Obbard *et al.* 2006; Palmer *et al.* 2018). To test whether the siRNA pathway is functional in hybrids between *D. melanogaster* and its sister species, we tested for the efficacy of RNAi in hybrids using a knockdown construct that targets the X-linked *white* gene (Lee *et al.* 2004). When the *white* gene is knocked down in flies, the eye color changes from the wild type red to white. Incomplete knockdown of this gene manifests as an intermediate color, which can be quantified. We generated hybrids between *D. melanogaster* females carrying this knockdown constructs and males from *D. simulans*, *D. mauritiana* and *D. sechellia* and measured the intensity of pigment as a readout of RNAi efficacy. Although the intensity of pigment was slightly greater in all hybrids than in comparable *D. melanogaster* genotypes, the reduction in eye pigmentation was not significantly different across hybrid genotypes (Figure 3). These results indicate that despite the rapid divergence of the genes involved in the siRNA pathway, this pathway remains functional in inter-species hybrids.

**Figure 3.**
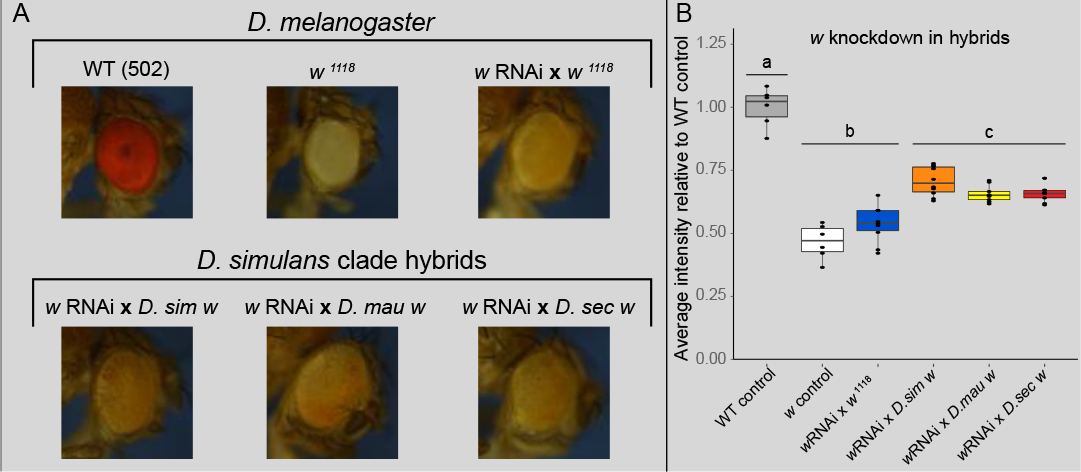
RNAi machinery functions in *D. melanogaster* – *D. simulans* clade hybrids. (A) Example eyes from the genotypes tested for eye color intensity. (B) Quantification of eye color intensity in control and hybrid genotypes. The lettered bars indicate categories that were significantly different from each other (Pairwise Wilcoxon Rank Sum test, p < 0.05, n=6). Hybrid pigment intensity is significantly reduced by the RNAi construct, and no hybrid genotype was significantly different from any other.

To directly test whether the level of knockdown of *gfzf*^*sib*^ is comparable across all three crosses, we performed the *gfzf*-knockdown hybrid rescue crosses with the three species and measured RNA expression levels of *gfzf*. We measured the levels of *gfzf* transcript from the parental species by RT-qPCR using primers that amplify only *gfzf*^*mel*^ or *gfzf*^*sib*^ in the hybrid females. We found that expression of the *gfzf*^*sib*^ allele is reduced in hybrids with all three species, and there is no significant difference between the magnitude of the reduction in any of the three species (Figure 4). Together these results indicate that the lack of rescue in *D. sechellia* is not due a failure to knock down the expression of *gfzf*^*sec*^.

**Figure 4.**
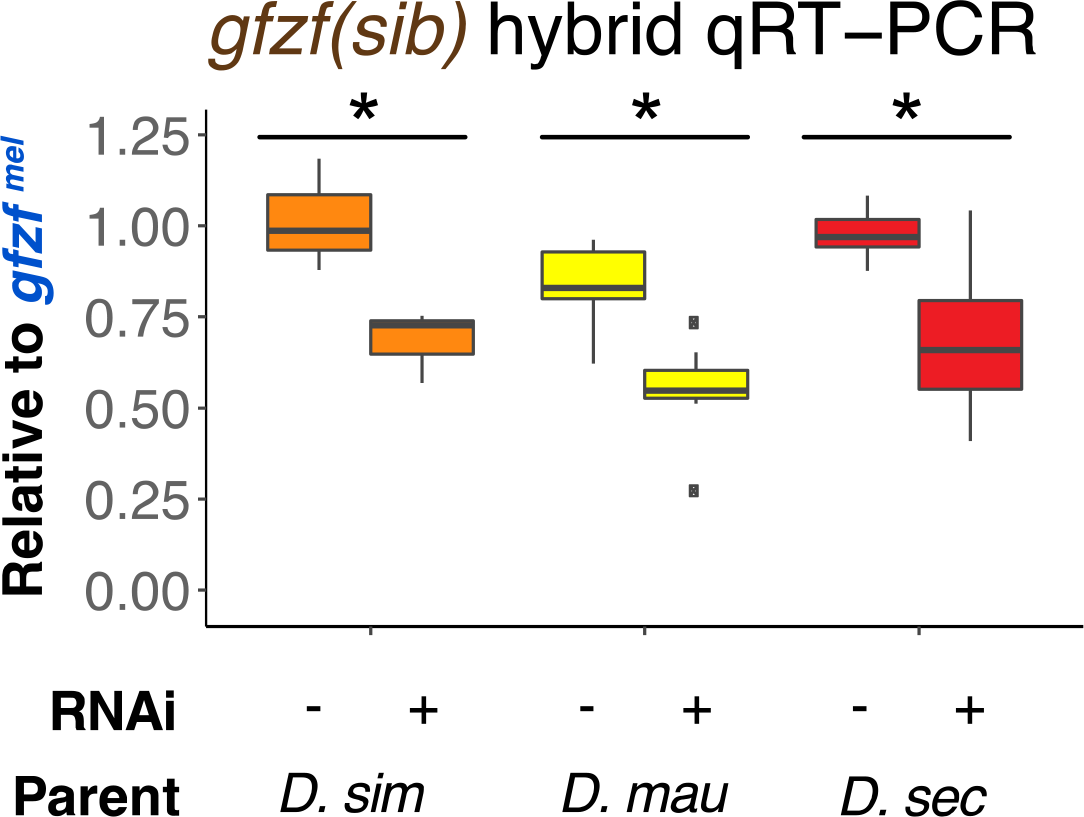
*gfzf*^*sib*^ RNAi construct reduces the expression of *gfzf*^*sib*^ in all three species. Expression of *gfzf* was normalized to *Rpl32* expression. Here values are presented as the ratio of *gfzf*^*sib*^ to *gfzf*^*mel*^. * p < 0.05 by Pairwise Wilcoxon Rank Sum test.

### *D. sechellia* has two dominant loci that confer resistance to hybrid male rescue

Our attempts to rescue *D. sechellia* hybrid males indicate that additional loci or known hybrid incompatibility loci fixed in the *D. sechellia* lineage to increase the strength of the hybrid incompatibility. We reasoned that as *D. sechellia* is resistant to hybrid male rescue, whereas *D. simulans* is not, recombinant genotypes between the two species would allow us to map the loci responsible for this trait. However, conventional multi-generation recombinant mapping schemes are untenable for this trait because the final cross must include a *D. melanogaster* female crossed to a *D. simulans-D. sechellia* hybrid male. Hybrid F1 males between *D. simulans* and *D. sechellia* are completely sterile, and thus any subsequent generations of recombinant males that could produce progeny with *D. melanogaster* would be biased by hybrid male sterility.

To circumvent this problem, we used a *D. simulans* attached-X (C(1)) stock to alter the direction of the *D. melanogaster* / *D. simulans-D. sechellia* cross while still preserving the genotype that we aimed to study (Figure 5A). The C(1) F1 *D. simulans* / *D. sechellia* hybrid females produce gametes that are recombinant for their autosomes, and either contain the *D. simulans* C(1) or a *D. sechellia* Y. When these hybrid females are crossed with a *D. melanogaster* male, this generates F1 triple-hybrid females with a *D. simulans* C(X) and F1 triple-hybrid males with a *D. melanogaster* X and a *D. sechellia* Y. This direction of the cross is susceptible to the cytoplasmic - nuclear incompatibly of a standard *D. melanogaster* male to *D. simulans* female cross, since the maternal factor from *D. simulans* (*Mhr*) interacts with the X chromosome from *D. melanogaster*, both of which are present in this cross. To remedy this, we recombined our RNAi- *gfzf*^*sib*^ transgene onto the *Zhr*^*1*^ chromosome, which is known to rescue the cytoplasmic - nuclear incompatibly (Sawamura and Yamamoto 1993). These genotypes allowed us to generate large numbers of triple-hybrid recombinant males, that when recovered should be enriched for *D. simulans* alleles that allow for their viability.

**Figure 5.**
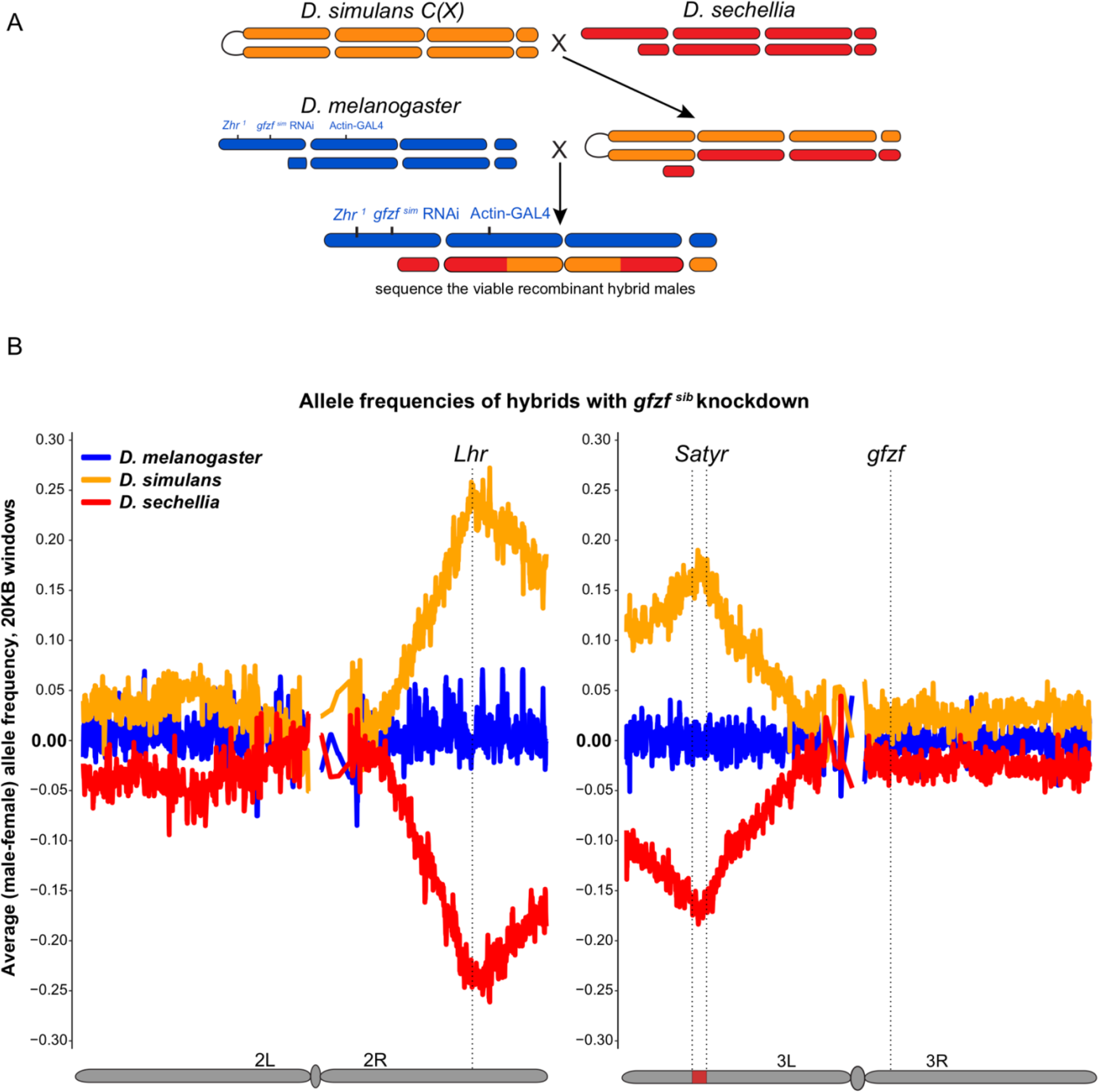
*D. sechellia* is resistant to hybrid male rescue by *gfzf*^*sib*^ RNAi due to two loci. (A) Cross for generating tri-hybrid progeny for mapping samples. Males and females were collected in three independent replicates for pooled genome sequencing. (B) Map of allele frequencies in the tri-hybrid males. For each sample, allele frequencies were calculated in 20KB windows. They were then normalized by subtracting the allele frequency for females in the same window, and outliers removed. This plot contains the average of all three replicates.

We used this crossing scheme to produce pools of triple-hybrid males and matched triple-hybrid females, starting from three inbred lines of *D. sechellia*. We then performed pooled whole genome sequencing on the males and the females from each replicate to measure the allele frequency of *D. simulans*, *D. sechellia*, and *D. melanogaster* alleles. In these experiments, the triple-hybrid females serve as a control for general effects on hybrid viability. When we analyzed the allele frequency of *D. simulans* and *D. sechellia* alleles in our samples, we found a striking result. There are two locations in the *D. simulans* genome that are highly enriched in viable hybrid males as opposed to the female samples (Figure 5B). The first peak of enrichment falls between 17.32MB and 17.50MB on chromosome 2R (*D. melanogaster* coordinates). This peak sits directly on top of *Lhr* (17.43MB), a known hybrid incompatibility gene in this system. The second peak appears on chromosome 3L between 8.62MB and 8.68MB (*D. melanogaster* coordinates). No genes near this region have previously been implicated in the *D. melanogaster / D. simulans* hybrid incompatibility. We name this locus *Sechellia aversion to hybrid rescue (Satyr)*.

Our data indicates that every male we recovered contained the *D. simulans Lhr* allele, while most males that we recovered contained the *Satyr* locus. Therefore, it appears that either *Lhr*^*sec*^ or *Satyr*^*sec*^ can prevent hybrid male rescue by *gfzf*^*sib*^ knockdown. Across the rest of the genome, there does appear to be a slight elevation in the recovery of *D. simulans* alleles, even in regions unlinked to *Lhr* or *Satyr*. The basis of this elevation is unclear – it may be due to the broad distribution of small effect genes, or due to some feature of genomic architecture that is present in one species but not the other. Importantly, we do not observe a peak in *D. simulans* allele frequency near *gfzf*, indicating that the *gfzf*^*sim*^ and *gfzf*^*sec*^ allele are equivalent in our experiment. It appears that the genomic architecture of this trait is dominated by two dominant large effect loci – *Lhr* and *Satyr*.

## Discussion

Our experiments are the first effort to map the variation of hybrid male rescue in the *D. simulans* clade by utilizing triple-hybrid crosses. Our data demonstrates that the interspecies variation in hybrid male rescue between *D. melanogaster* and the *D. simulans* clade is not dispersed broadly across the genome, but is instead confined to just two major effect loci. In the hybrid, these loci are still present with the equivalent alleles from *D. melanogaster*, meaning that their effect on hybrid male rescue is dominant. Since we find these differences to be fixed between the three species of the *D. simulans* clade, it is likely that *D. sechellia* became fixed for these changes after its split from *D. simulans* and *D. mauritiana*. Though these are young species of *Drosophila*, they have undergone significant divergence from one another – even though all three species produce sterile males when hybridized with each other, the nature of that male sterility depends on the parents involved (Zeng and Singh 1993).

The association with changes in *Lhr* in this genetic map imply that changes at *Lhr* have risen to fixation in *D. sechellia*. It is also possible that island colonization and subsequent bottleneck in effective population size during the establishment of *D. sechellia* as an island species cause the fixation of variants in *Lhr.* However, as *Lhr* appears to play a key role in the suppression of transposable elements, it is possible that an arms race with selfish elements of the genome have placed *Lhr* under strong selective pressure post *D. simulans* – *D. sechellia* speciation. This raises the intriguing possibility that *Lhr*^*sim*^ and *Lhr*^*sec*^ may have different capabilities to regulate TEs in a *D. simulans* background. Further, understanding the difference between the two forms of *Lhr* might link directly to its role in the hybrid incompatibility between *D. melanogaster* and the *D. simulans* clade.

Additionally, our data suggests that variants at *Lhr* have an effect on rescue by reduction of *gfzf*^*sib*^. This adds a much more direct connection between *Lhr* and *gfzf* than has previously been identified. Combined with our recent work to show that *Hmr* and *gfzf* co-localize on DNA in hybrids (Cooper *et al.* 2018), a new picture of these three hybrid incompatibility genes is beginning to emerge. Rather than acting as additive entities, it appears that all three of these genes have the ability to influence the actions of each other. It is still unclear what the direct molecular mechanism of interaction for all three of these genes may be, but our data support that they act to influence the same molecular property rather than working as additive effects.

We have also uncovered a new locus, *Satyr*, that acts as an additional dominant hybrid incompatibility locus between *D. melanogaster* and *D. sechellia.* This locus resides near 8.6MB (*D. melanogaster* coordinates) on chromosome 3L. One intriguing possibility is that the action of this locus is not confined to the *D. melanogaster* – *D. sechellia* hybridization, and that loss of function mutations at this locus from any of the *D. simulans* sibling species may be sufficient to rescue hybrid males. Importantly, our approach relies on sensitization to rescue by reducing rather than completely removing *gfzf*^*sec*^. Therefore it is possible that complete loss of function at *gfzf*^*sec*^, *Lhr*^*sec*^, or *Satyr*^*sec*^ would be capable of full rescue of hybrid males.

In our experiments, we used RNAi to reduce the expression of *gfzf*^*sib*^ to rescue hybrid F1 males. This likely has two major implications for our results. First, in our system we know that rescue was not complete in *D. simulans* (near 68%), indicated that we were much closer to a sensitizing threshold for rescue than might be obtained by genetic ablation of *Lhr* or *gfzf*. This means our system may have allowed us to detect loci that would have otherwise been missed by a strong rescue allele. Second, our conclusions cannot make statements about the sufficiency of completely missing these loci as a means of rescuing hybrid males. However, our results are still clear on the point that *Lhr*^*sec*^ is a stronger hybrid incompatibility allele than *Lhr*^*sim*^, even if complete loss of either would rescue hybrid F1 males. Importantly, our results indicate that *gfzf*^*sec*^ is not a stronger hybrid incompatibility allele than *gfzf*^*sim*^, since there is no elevation of the *D. simulans* allele frequency at *gfzf* in our data.

Previous efforts to locate hybrid incompatibility loci have relied on the recovery of natural alleles (Watanabe 1979; Hutter and Ashburner 1987), deficiency screens from *D. melanogaster* (Presgraves 2003; Cuykendall *et al.* 2014), and a mutagenesis screen in *D. simulans* (Phadnis *et al.* 2015). In all of these instances it is entirely plausible that a dominant hybrid incompatibility at the *Satyr* locus would have been missed – the mutagenesis screen was not to saturation, as no alleles at *Lhr* were recovered. Our work highlights the important role that interspecies variation can play as a general tool for dissecting hybrid incompatibilities (Orr and Coyne 1989), and demonstrates the continued evolution of the *D. melanogaster – D. simulans* clade hybrid incompatibility through a few large effect loci in the *D. sechellia* genome.

## Materials and Methods

### Fly strains

The details natural populations and species variants that we acquired for this experiment can be found in Table 1. These lines were gifts from H.S. Malik, D. Matute, or acquired from the Drosophila Species Stock Center. For our triple-hybrid mapping cross, we generated several lines. First, we build a recombinant RNAi- *gfzf*^*sib*^, *Zhr*^*1*^ chromosome by recovering the products of RNAi- *gfzf*^*sib*^ (Phadnis *et al.* 2015) crossed to *Zhr*^*1*^ (Bloomington Stock Center 25140) over an *FM7i* balancer. We then confirmed the ability of this chromosome to rescue hybrid F1 males by crossing it to the C(X) *D. simulans* stock (Sawamura *et al.* 1993). Next, we generated three independent stocks of *D. sechellia w* (Drosophila Species Stock Center 14021-0248.15) by single pair inbreeding three replicates of the base stock for five generations. To induce our RNAi system, we crossed the RNAi- *gfzf*^*sib*^, *Zhr*^*1*^ chromosome to an Actin5C-GAL4 / CyO line (Bloomington Stock Center 25374), and crossed the resulting CyO^+^ F1 males to C(X) *D. simulans / D.sechellia* F1 hybrid females to make the triple-hybrid progeny.

**Table 1.** Species and Strains.

| Name | Species | Origin |
| --- | --- | --- |
| WT (07) | *D. simulans* | Wanie-Rukula, Congo |
| WT (08) | *D. simulans* | Wanie-Rukula, Congo |
| *w<sup>501</sup>* | *D. simulans* | Drosophila Species Stock Center |
| C(X) | *D. simulans* | Drosophila Species Stock Center |
| iso-105 | *D. mauritiana* | Drosophila Species Stock Center |
| w139 | *D. mauritiana* | Malik Lab |
| w140 | *D. mauritiana* | Malik Lab |
| WT(09) | *D. mauritiana* | Malik Lab |
| iso-75 | *D. mauritiana* | Malik Lab |
| *w[1]* | *D. mauritiana* | Drosophila Species Stock Center |
| *w* | *D. sechellia* | Drosophila Species Stock Center |
| WT(03) | *D. sechellia* | Drosophila Species Stock Center |
| NF13 | *D. sechellia* | Matute Lab |
| NF14 | *D. sechellia* | Matute Lab |
| ArvoB3 | *D. sechellia* | Matute Lab |

### Fly husbandry

For our initial tests of hybrid male rescue, we allowed parental flies to mate for 2 days at 25C before flipping them to fresh media. We incubated the vials containing hybrid progeny at 18C, as during the larval stages hybrid larvae become extremely temperature sensitive (Barbash *et al.* 2000). We counted the progeny at 23 days post mating. In generating the triple-hybrid flies, we used a different mating scheme as we found that the C(X) genotype has high rates of lethality at 25C. For these crosses, we allowed mating at 21C for 2 days, followed by incubating the progeny at 18C until 23 days post mating.

### Measuring *gfzf* expression in hybrids

To determine the genotypes of our samples, we removed the head and probed for the presence of the RNAi construct and GAL4 by PCR. To measure *gfzf* expression, we extracted RNA from the remainder of the body using the DirectZol RNA Miniprep Kit (Zymo Research) and generated cDNA using SuperScript III (Thermo Fisher Scientific). For RT-qPCR, we used iTaq Syber Green (BioRad). We measured the abundance of *gfzf*^*mel*^, *gfzf*^*sib*^, and Rpl32 as a loading control using the following primers: *gfzf* F(both species): CCGGACATGGACCTCTCAAA, *gfzf* R (mel): GGGACACGGATAATGATGCAG, *gfzf* R (sim): CTTTGGGACACGGATCTGCT, RPL32 F: ATGCTAAGCTGTCGCACAAATG, R: GTTCGATCCGTAACCGATGT. We rejected any samples in which our no-RT controls showed signs of amplification. To compare expression levels, we first normalized both *gfzf* samples to the Rpl32 control, and then determined the ratio of *gfzf*^*sib*^ to *gfzf*^*mel*^ expression. We checked for statistical significance in our samples using a Pairwise Wilcoxon Rank Sum test in R.

### Measuring eye pigment in *w*-RNAi

To measure the pigment intensity of eyes in our different genotypes, we gathered images of both eyes from individual flies using a Lieca MC120 HD camera on a Lieca MC165 FC dissection scope with overhead illumination. To control for changes in ambient lighting, we included a piece of blue construction paper as the background, and made sure to capture the image such that segments of the construction paper were not in the shadow of the fly. We used the gray scale of these images to measure pixel intensity in ImageJ, and normalized the values to that of the construction paper in the background. We normalized all values to the mean of the WT control, and checked for a statistical difference between the samples using a Pairwise Wilcoxon Rank Sum test in R.

### Whole fly DNA extraction for pooled genome sequencing

To extract DNA for whole genome sequencing, we used the DNeasy Blood and Tissue kit (Qiagen). We pooled our 350 triple-hybrids by simultaneously by freezing all samples in liquid nitrogen and grinding them together with a mortar and pestle, and immediately using the frozen ground tissue as the input for the DNeasy kit. We repeated this process for each of the triple-hybrid male and paired triple-hybrid female samples. For the parental lines that we sequenced, we extracted DNA from a pool of 50 flies, half male and half female.

### Pooled whole genome sequencing

To measure allele frequencies in our triple-hybrid samples, we used the PCR-free Illumina Novaseq platform to generate paired end reads of the pooled sample. To generate accurate calls of variants in our different lines, we sequenced all six of our parental lines using the Hi-Seq Illumina platform. Library prep and sequencing was carried out by the Huntsman Cancer Institute High-Throughput Genomics and Bioinformatics Analysis Shared Resource.

### Sequence alignment and allele frequency analysis

We trimmed sequencing reads for quality using PicardTools. We aligned the reads to the *D. melanogaster* reference genome (r6.24 at the time of analysis) using bwa (Li and Durbin 2009). We called variants and re-aligned reads based on these variant calls using GATK 3.6 (McKenna *et al.* 2010). To find positions that would allow us to measure allele frequency in the three species, we wrote our own code to parse vcf files and identify tri-partite SNP positions (for our analyses, we did not use indels as tri-partite SNPs were at high enough frequency). To analyze allele frequencies, we scanned the genome in 20KB windows and measured the relative abundance of *D. melanogaster*, *D. simulans*, and *D. sechellia* SNPs from high quality sites. We paired these windows between male and female samples, calculated the difference in allele frequency between males and females for all three of the parental SNP types. The plot that we report in Figure 2 is the average allele frequency in each window for all three replicates. All of our code can be found at github.com/jcooper036/tri_hybid_mapping.

## Data Accessibility

All of the genomic sequencing data for this project is available on the Sequence Read Archive accession number SRP190327. It can also be accessed via the BioProject accession number PRJNA530263.

## Acknowledgements

We thank the labs of H. S. Malik and D. Matute for sharing their fly lines with us. We thank S. Phadnis for her continued support. Research reported in this publication utilized the High-Throughput Genomics and Bioinformatic Analysis Shared Resource at Huntsman Cancer Institute at the University of Utah and was supported by the National Cancer Institute of the National Institutes of Health under Award Number P30CA042014. The content is solely the responsibility of the authors and does not necessarily represent the official views of the NIH.

